# C/EBPδ-induced epigenetic changes control the dynamic gene transcription of S100A8 and S100A9

**DOI:** 10.1101/2021.12.05.471309

**Authors:** Saskia-L. Jauch-Speer, Jonas Wolf, Marisol Herrera-Rivero, Leonie Martens, Achmet Imam Chasan, Anika Witten, Birgit Markus, Bernhard Schieffer, Thomas Vogl, Monika Stoll, Johannes Roth, Olesja Fehler

## Abstract

The proinflammatory alarmins S100A8 and S100A9 are among the most abundant proteins in neutrophils and monocytes but completely silenced after differentiation to macrophages. The molecular mechanisms of the extraordinarily dynamic transcriptional regulation of *s100a8* and *s100a9* genes, however, are only barely understood. Using an unbiased genome-wide CRISPR/Cas9 knockout based screening approach in immortalized murine monocytes we identified the transcription factor C/EBPδ as a central regulator of S100A8 and S100A9 expression. *S100a8* and *S100a9* expression was further controlled by the C/EBPδ-antagonists ATF3 and FBXW7. We confirmed the clinical relevance of this regulatory network in subpopulations of human monocytes in a clinical cohort of cardiovascular patients. Moreover, we identified specific C/EBPδ-binding sites within *s100a8* and *s100a9* promoter regions, and demonstrated that C/EBPδ-dependent JMJD3-mediated demethylation of H3K27me_3_ is indispensable for their expression. Overall, our work uncovered C/EBPδ as a novel regulator of S100A8 and S100A9 expression. Therefore, C/EBPδ represents a promising target for modulation of inflammatory conditions that are characterised by S100A8 and S100A9 overexpression.

## INTRODUCTION

As the first line of immune defense, both, monocytes and neutrophils are important for the modulation of the innate immune response. To amplify the immune response at sites of inflammation, the activation of further immune cells is required, mediated by the release of signaling molecules such as chemokines and DAMPs (damage-associated molecular patterns). The two members of the S100 family, S100A8 and S100A9, also termed myeloid-related proteins 8 and 14 (MRP8 and MRP14), respectively, belong to the group of DAMPs or so-called alarmins. Their primary expression is referred to myeloid-lineage derived cells, particularly neutrophils and monocytes, where S100A8 and S100A9 are predominantly present as a heterodimeric complex, also called calprotectin (Austermann, Spiekermann and Roth, 2018).

Intracellularly, S100A8/S100A9 complexes represent up to 40% of the soluble protein content in neutrophils and about 5% in monocytes (Hessian, Edgeworth and Hogg, 1993). However, in mature macrophages, protein and mRNA expression of these factors is completely downregulated. This data indicates that expression of S100A8 and S100A9 is controlled by the most dynamic promotors in myeloid cells. The S100A8/A9 complex interacts with the cytoskeleton in a calcium-dependent manner. Calcium-induced (S100A8/A9)_2_ tetramer promotes tubulin polymerization and microtubule bundling, thereby affecting transendothelial migration of phagocytes (Leukert *et al.*, 2006). During inflammation or tissue damage S100A8/A9 is actively secreted by neutrophils and monocytes, and represents the most abundant DAMP/alarmin activating inflammatory processes in infection, cancer, autoimmunity and cardiovascular diseases. The S100A8/A9 complex is recognized by Toll-like receptor 4 (TLR4), which leads to the production of proinflammatory cytokines and chemokines (Fassl *et al.*, 2015). Accordingly, S100A9 KO mice exhibit decreased pathogenic outcomes in several mouse models of disease, such as sepsis (Vogl *et al.*, 2007), autoimmune disease (Loser *et al.*, 2010) or arthritis (Van Lent *et al.*, 2008). In addition, S100A8 and S100A9 are highly abundant during infectious diseases and exhibit anti-microbial activities. The S100A8/A9 complex plays a crucial role in host defense against bacterial and fungal pathogens by sequestering manganese and zinc ions which compete with high affinity bacterial transporters to import these essential nutrient metals (Kehl-Fie and Skaar, 2010; Kehl-Fie *et al.*, 2011). In contrast to the proinflammatory role of S100A8/A9, also regulatory functions in terms of hyporesponsiveness in phagocytes, resembling a classical endotoxin-induced tolerance, have been described (Freise *et al.*, 2019).

In humans, S100A8/A9 is the most abundant alarmin in many clinically relevant diseases, and is closely associated with disease activity in rheumatoid arthritis, inflammatory bowel disease, sepsis, cardiovascular diseases, multiple sclerosis, acute lung injury and psoriasis (Foell *et al.*, 2004). Altered S100A8/A9 expression has also been found in different cancer types, including gastric, colorectal, breast, lung, prostate and liver cancer (Cross *et al.*, 2005). Despite the high expression in neutrophils and monocytes under inflammatory conditions, and the strong effects of S100A8 and S100A9 on disease activities, transcriptional mechanisms regulating these extreme dynamics of gene expression remain unclear. Identifying the mechanisms regulating S100A8 and S100A9 gene expression may open new insights into the pathological processes involving S100A8/A9 during inflammatory conditions.

So far, several potential transcription factors modulating S100A8 and S100A9 expression have been described (Kuruto-Niwa *et al.*, 1998; Fujiu, Manabe and Nagai, 2011; Lee *et al.*, 2012; Liu *et al.*, 2016; Yang *et al.*, 2017), but their functional relevance remains unresolved. Many of the stated studies used malignant immortalized cell lines or even cell models whose homologous primary cells do not express these genes at all.

To overcame difficulties of artificial expression and malignant cell lines we used ER (estrogen-regulated) Hoxb8 cells, estrogen dependent transiently immortalized myeloid precursor cells that can be differentiated to primary monocytes and granulocytes upon estrogen-withdrawal (G. G. Wang *et al.*, 2006), and show the physiologically high dynamics of S100A8 and S100A9 mRNA and protein expression during differentiation. In order to detect genes involved in the regulation of S100A8/A9 expression in an unbiased manner, we used a mouse Genome-Scale CRISPR/Cas9 Knockout (GeCKO) library and screened for monocytes with reduced or absent S100A9 expression. We thereby identified the CCAAT/enhancer binding protein-family member C/EBPδ as a direct transcriptional regulator of S100A8/A9. Furthermore, we found that the epigenetic factor JMJD3 contributes to S100A8 and A9 expression in monocytes by erasure of the repressive histone mark H3K27me_3_ at *s100a8* and *a9* promoter regions. Moreover, we confirmed the clinical relevance of this network in specific monocyte subpopulations in a cohort of patients with cardiovascular disease.

## RESULTS

### Genome-wide CRISPR/Cas9 Knockout screen reveals C/EBPδ as a regulatory factor of S100A9 expression

To detect genes involved in the regulation of S100A8/A9 expression, we established the mouse GeCKO (Genome-Scale CRISPR-Cas9 Knockout) lentiviral pooled library designed in Cas9 expressing ER-Hoxb8 cells. The used library contained a large mixture of CRISPR sgRNA constructs, where six gRNAs per target gene increase efficiency and enable the analysis of the molecular effects of many thousand genes in one experiment. After infection of Cas9 expressing ER-Hoxb8 precursor cells with CRISPR library lentiviral particles, the cells were differentiated for 3 days in the presence of GM-CSF to induce S100A8 and S100A9 expression. Because we hypothesized that the parallel S100A8 and S100A9 expression is based on a common regulatory mechanism, we assumed that screening of one of the two alarmins is sufficient in the first step. Therefore, cells with no or low S100A9 expression were selected by sorting and considered as hits, whereas the remaining cells functioned as reference cells. To exclude phenotypes that are S100A9^low/neg^ due to differentiation defects, we pre-gated for CD11b^+^Ly-6C^+^ monocytes. DNA of sorted cell pools was purified and analysed by NGS (Figure 1, A). Intracellular S100A9-FITC staining of precursor and differentiated Cas9 ER-Hoxb8 control monocytes was used as standard for definition of sorting gates. Differentiated Cas9-library ER-Hoxb8 monocytes showed a wider distribution among the gates, indicating the presence of S100A9^low/neg^ expressing cells due to disruptions of regulatory genes caused by CRISPR/Cas9. A small amount of S100A9^neg^ sorted cells were re-analysed by immunoblotting to validate S100A9 deficiency in this cell population (Figure 1, B). Analysis of CRISPR KO library screen using the MAGeCK method(Li *et al.*, 2014) resulted in a list of genes for which the respective gRNAs were enriched in the hits sample. The highest number of gRNAs found within the top 20 hits targeted *cebpd*, a gene encoding for a member of the CCAAT/Enhancer-Binding-protein family, C/EBPδ (Figure 1, C).

**Figure 1:**
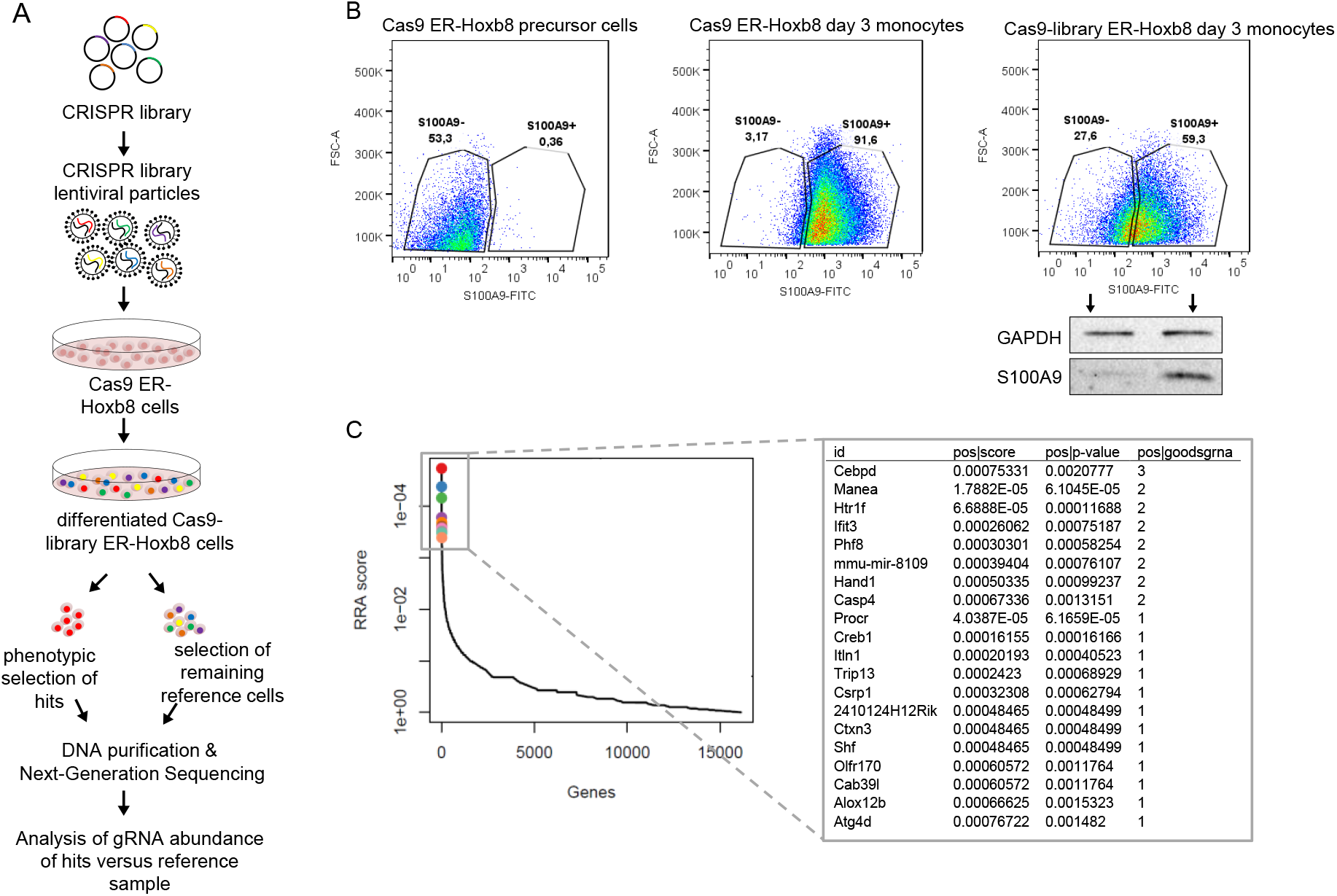
Genome-Scale CRISPR Knockout lentiviral pooled library screen to identify S100A9-regulators. (A) For genome-wide screen, over 100,000 plasmids, each containing a guide RNA towards different early consecutive exons, were packaged into lentiviral particles. Cas9 expressing ER-Hoxb8 cells were pool-transduced, selected and differentiated to induce S100A9 expression. Hits and reference cells were collected by sorting according to their phenotypes of interest. DNA of both samples was purified for next-generation sequencing and subsequent analysis. (B) Precursor and differentiated Cas9 and Cas9-library ER-Hoxb8 cells were stained intracellularly for S100A9 using a FITC-labelled antibody. Cas9-library ER-Hoxb8 day 3 monocytes with no or lower S100A9 expression were sorted as hits, the remaining cells served as reference cells. (C) Data was analysed using the MAGeCK software for identification of enriched guide RNAs in the hit sample. Corresponding genes were rank-ordered by robust rank aggregation (RRA) scores. The list states the top 20 genes according to RRA scores, arranged after the number of guides that are enriched in the hit sample

### Decreased s100a8 and s100a9 expression in C/EBPδ KO monocytes

We confirmed extraordinarily high dynamics of S100A8 and S100A9 expression during monocyte/macrophage differentiation. ER-Hoxb8-derived monocytes show an about 590-fold increase in *s100a8* mRNA expression and an about 1,800-fold increase of *s100a9* mRNA expression on day 2 compared to day 0 of differentiation. At day 5 the *s100a8* mRNA expression is already about 70-fold, the *s100a9* mRNA expression about 110-fold down-regulated compared to day 2 of differentiation (Figure 2 A and B).

**Figure 2:**
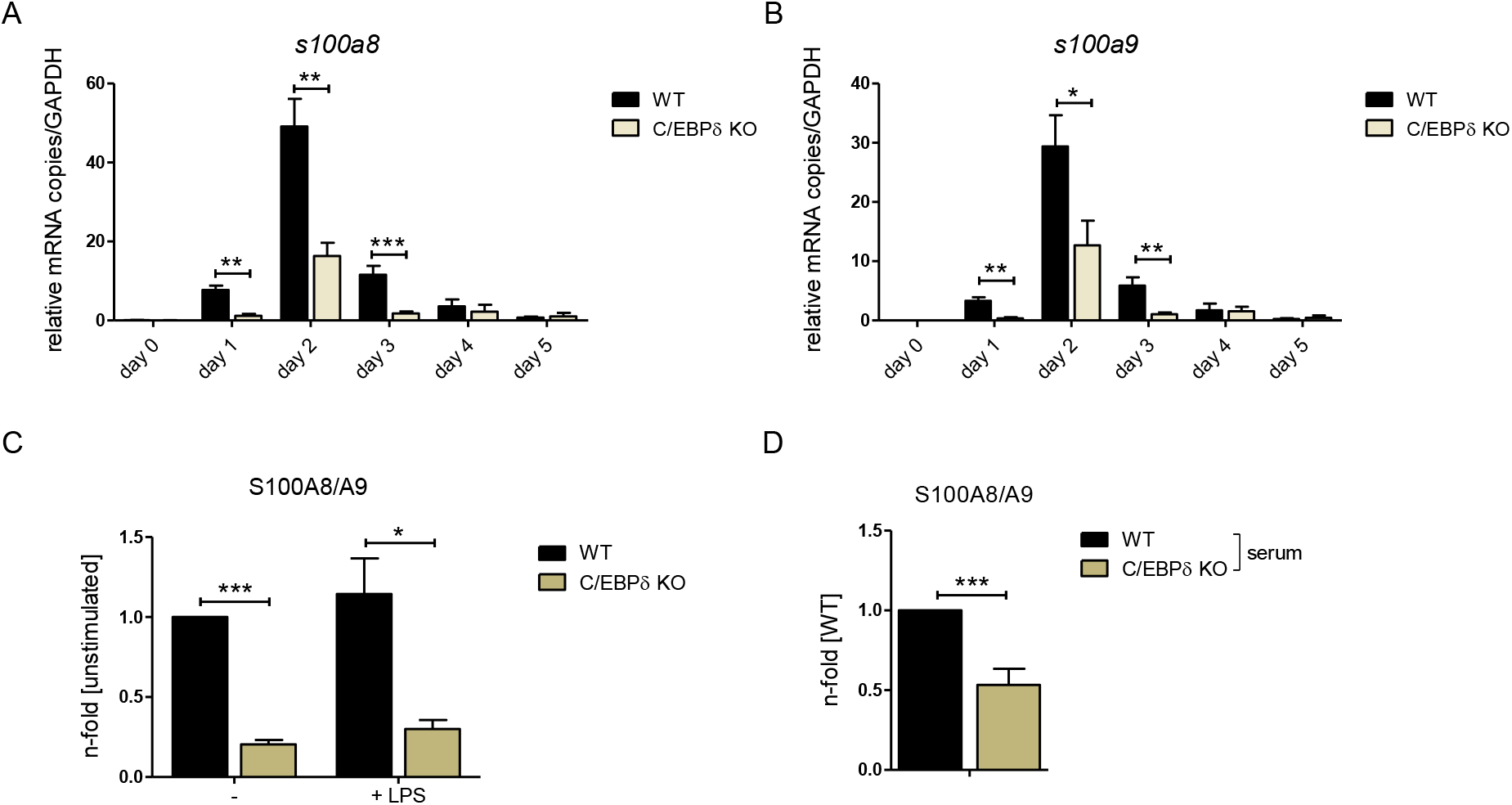
Figure 3: S100A8 and S100A9 expression in WT and C/EBPδ KO ER-Hoxb8 monocytes. (A) Relative s100a8 and (B) s100a9 mRNA levels were measured using qRT-PCR (n = 3-8). (C) S100A8/A9 concentrations in supernatant of differentiation day 4 of WT and C/EBPδ KO monocytes stimulated with 10 ng LPS for 4 hours or left untreated (n = 3) and (D) serum concentrations of S100A8/A9 in WT and C/EBPδ KO mice were quantified using our in-house mouse S100A8/S100A9 sandwich ELISA (n = 6). Values are the means ± SEM. *P < 0.05, **P < 0.01, ***P < 0.001, by two-tailed Student’s t test. See also Figure supplements 1, 2, and 3.

We confirmed the relevance of C/EBPδ for S100-expression by creating independent C/EBPδ-deficient ER-Hoxb8 cells from C/EBPδ KO mice. Not only on differentiation day 3, but already at the very beginning of differentiation, when *s100a8* and *a9* levels start to rise, C/EBPδ-deficient ER-Hoxb8 monocytes showed significantly reduced levels of both, *s100a8* (Figure 2, A) and *s100a9* mRNAs (Figure 2, B), compared to WT controls. The same effect was detectable in C/EBPδ-deficient ER-Hoxb8 cells that were differentiated into the neutrophilic lineage (Figure 2 – figure supplement 1, A). Accordingly, *cebpd* and *s100a8* and *a9* mRNAs were co-expressed in differentiating WT monocytes and neutrophils, supporting a mechanistic connection (Figure 2 – figure supplement 1, B and C). WT monocytes secret significant S100A8/A9-protein amounts, whereas the supernatant of C/EBPδ KO monocytes has up to 80% less S100A8/A9 (Figure 2, C). Consistent with this, serum S100A8/A9-levels are significantly decreased in C/EBPδ KO mice (Figure 2, D). Although the proinflammatory molecule S100A8/A9 is strongly reduced in the C/EBPδ KO monocytes, these cells exhibit no general alterations of inflammatory functions, indicating a rather specific effect on S100A8/A9-regulation than a general attenuation of inflammatory signaling due to C/EBPδ-deficiency. Phagocytosis capacities are even elevated in C/EBPδ KO monocytes, very likely through an already known PTX3-dependent mechanism (Ko *et al.*, 2012) (Figure 2 – figure supplement 2, A and B), whereas ROS production is not influenced by C/EBPδ deficiency (Figure 2 – figure supplement 2, C).

Interestingly, none of the transcription factors previously reported to target S100A8 and A9 were found within the hit list of our CRISPR KO library screen (Supplementary Table S1). Nevertheless, to test the published candidate transcription factors ATF3, STAT3, KLF5, IRF7 and C/EBPβ for their effects on S100A8 and A9-regulation, we created single KO ER-Hoxb8 cell lines of each individual candidate transcription factor. At no-time point during differentiation, deficiency of the stated candidate transcription factors affected *s100a8* and *s1009* expression, whereas C/EBPδ-deficiency has a strong attenuating effect on *s100a8* and *s1009* expression, as shown on day 2 (Figure 2 – figure supplement 3, A and B).

### Enhanced C/EBPδ expression induces S100A8 and S100A9 expression

To test the impact of C/EBPδ-induction on S100-alarmin expression, we infected C/EBPδ-deficient ER-Hoxb8 cells with lentiviral particles carrying a tet-on system for doxycycline-inducible 3xFlag-C/EBPδ expression (Figure 3, A). Doxycycline treatment led to expression of the fusion protein 3xFlag-C/EBPδ, as revealed by western blot analysis (Figure 3, B) and by qRT-PCR in comparison to C/EBPδ-deficient cells (Figure 3, C). Induction of 3xFlag-C/EBPδ upon doxycycline-treatment led to increased *s100a8* and *s100a9* mRNA levels. C*ebpd*, *s100a8* and *s100a9* mRNA levels in doxycycline-treated TRE_3xFlag-C/EBPδ cells were comparable to WT cells at the same differentiation stage (Figure 3, D), demonstrating a positive effect of C/EBPδ expression on S100A8/A9 regulation. Knockout of ATF3, a known regulatory attenuator of *cebpd* expression (Litvak *et al.*, 2009), in ER-Hoxb8 monocytes led to S100A8/A9 overexpression (Figure 2 – figure supplement 3), especially during early stages of differentiation (Figure 3, E). ATF3 KO cells showed significantly elevated *cebpd* level, indicating a C/EBPδ-mediated effect on the expression of *s100a8* and *a9* (Figure 3, F). In the next step, we created FBXW7-deficient monocytes. FBXW7 is another well-known attenuator of C/EBPδ expression (Balamurugan *et al.*, 2013). Lack of this antagonist resulted in an even higher overexpression of *s100a8* and *s100a9* (Figure 3, G) with huge increases of *cebpd* levels (Figure 3, H).

**Figure 3:**
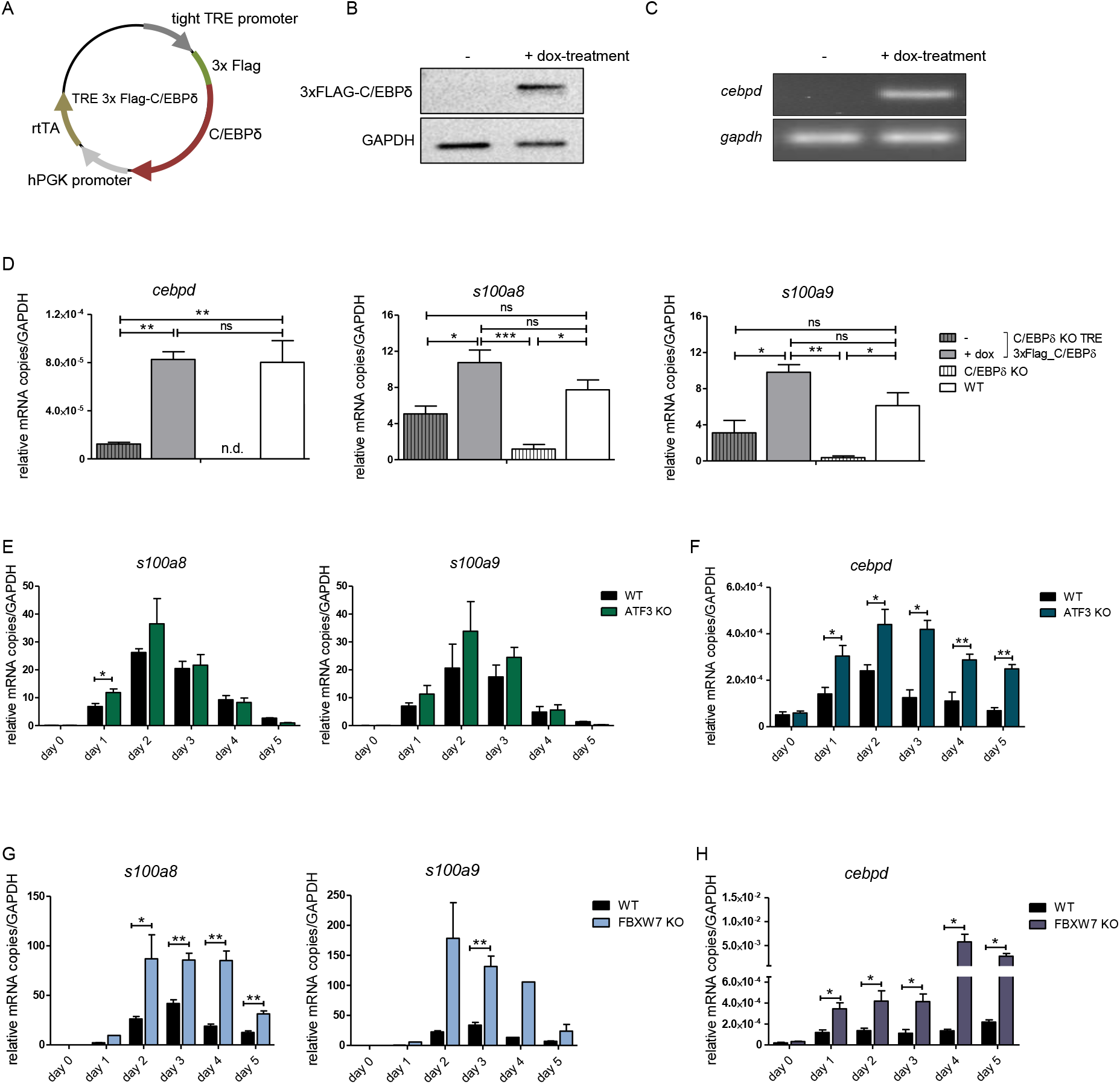
Figure 4: S100A8 and S100A9 expression in differentiated ER-Hoxb8 cells is dependent on C/EBPδ abundancy. (A) Tet-On construct of inducible 3xFlag-C/EBPδ expression due to constitutively expressed rtTA (reverse tetracycline-controlled transactivator) that binds to TRE promoter upon doxycycline-treatment was transduced into C/EBPδ KO ER-Hoxb8 cells. (B) Induction of 3xFlag-C/EBPδ upon doxycycline treatment (2 µg/mL, 24h) was analysed by western blot and (C) qRT-PCR in comparison to untreated cells. (D) Induction of 3xFlag-C/EBPδ was also analysed by qRT-PCR (cebpd), as well as expression of s100a8 and a9 mRNAs, in untreated and dox-treated C/EBPδ KO TRE_3xFlag-C/EBPδ monocytes and in comparison to WT and C/EBPδ KO monocytes on differentiation day 1 (n = 3). (E, G) S100a8 and s100a9, (F, H) and cebpd mRNA levels were measured using qRT-PCR in precursor and differentiated WT and ATF3 KO (E, F) and in WT and FBXW7 KO (G, H) ER-Hoxb8 monocytes (n = 3-4). Values are the means ± SEM. *P < 0.05, **P < 0.01, by one-way ANOVA with Bonferroni correction (D) and by two-tailed Student’s t test (E-H).

To confirm the biomedical relevance of the identified molecular network, we analysed the expression of these genes in PBMCs and monocyte subpopulations of a subset of participants in the BioNRW Study. Here, we found up-regulation of S100A8, S100A9 and CEBPD in PBMCs of sCAD/MI cases, compared against controls (Figure 4, A), together with a positive correlation of S100A8 and S100A9 expression with that of C/EBPD in these cells (Figure 4, B and C). Moreover, there was also significant up-regulation of these three genes specifically in classical monocytes, compared to intermediate and non-classical monocyte subpopulations (Figure 4, D and E, for further comparisons see Supplementary Table S9). A strong positive correlation between the expression of S100A8 and A9 and C/EBPD in these monocyte subpopulations was found, suggesting that the expression of these genes is mainly associated with the subset of pro-inflammatory monocytes. Interestingly, we also found significant albeit milder, negative correlations between the expression of C/EBPD and its antagonists FBXW7 and ATF3 in monocytes (Figure 4, F and G).

**Figure 4:**
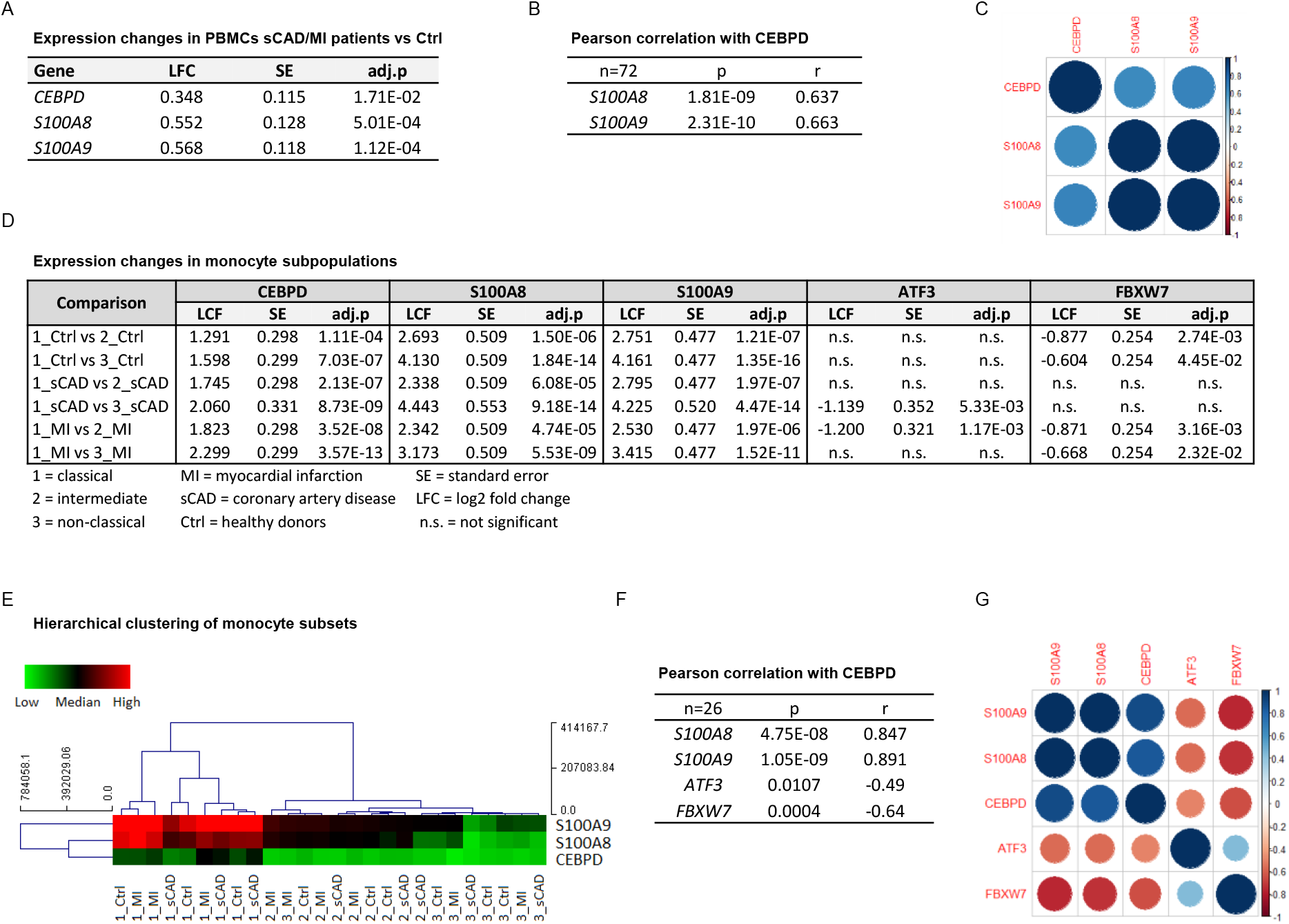
C/EBPδ expression positively correlates with S100A8 and S100A9 expression in proinflammatory monocytes of MI/sCAD patients. (A) Gene expression changes detected by RNA-seq in PBMCs of BioNRW participants (n=72, sCAD/MI vs Ctrl). LFC = log2 fold change, SE = standard error, and adj.p = adjusted P-value. (B) Pearson correlation coefficient = r, P-value = p in PBMCs and (C) corresponding correlation matrix. (D) Gene expression changes of CEBPD, S100A8, S100A9, ATF3 and FBXW7 detected by RNA-seq in monocyte subpopulations of BioNRW participants (n=26,from 3 individuals in each of the sCAD, MI and Ctrl diagnostic groups). (E) Hierarchical clustering of S100A8-, S100A9- and CEBPD normalised counts (using Euclidean distance metric with complete linkage). Shown are classical (1), intermediate (2) and non-classical (3) monocytes of healthy donors (Ctrl), myocardial infarction (MI) and stable coronary artery disease (sCAD) patients. (F) Pearson correlation coefficient = r, P-value = p in monocytes and (G) corresponding correlation matrix. See also Supplementary Table S9.

### C/EBPδ-binding sites within s100a8 and s100a9 promoter regions

Chromatin immunoprecipitation revealed 3xFlag-C/EBPδ binding on *s100a8* and *s100a9* promoter regions just before or within the predicted enhancers (Figure 5, A). Co-transfection of HEK293T cells with GFP expressing S100-reporter vectors, together with doxycycline-inducible 3xFlag-C/EBPδ vector (TRE_3xFlag-C/EBPδ) or its backbone lacking the 3xFlag-C/EBPδ construct (TRE_ctrl), was performed to further examine specific C/EBPδ-binding (Figure 5, B). Doxycycline-treatment resulted in 3xFlag-C/EBPδ protein expression after 24h post-transfection in 3xFlag-C/EBPδ vector transfected cells (Figure 5, C). Transfection of either *s100a8*-reporter construct (Figure 5, D) or *s100a9*-reporter construct (Figure 5, F) together with 3xFlag-C/EBPδ vector led to enhanced GFP expression upon doxycycline-treatment, in comparison to co-transfection with backbone plasmid (TRE_ctrl). Next, we modified predicted C/EBP-binding sites on S100 promoters by mutagenesis of the corresponding vectors. Again, co-transfection of mutated S100 reporter vectors and doxycycline-dependent 3xFlag-C/EBPδ vector was performed to analyse the relevance of specific C/EBPδ-binding sites. Two sites within the *s100a8* promoter region, stated as site 2 and site 3 (Figure 5, E), and one within the *s100a9* promoter region, stated as site 4 (Figure 5, G), caused a reduced or absent GFP expression upon co-transfection when deleted. These binding sites, in turn, were located within the *s100a8* and *s100a9* promoter regions where C/EBPδ-binding was confirmed by ChIP (Figure 5, A).

**Figure 5:**
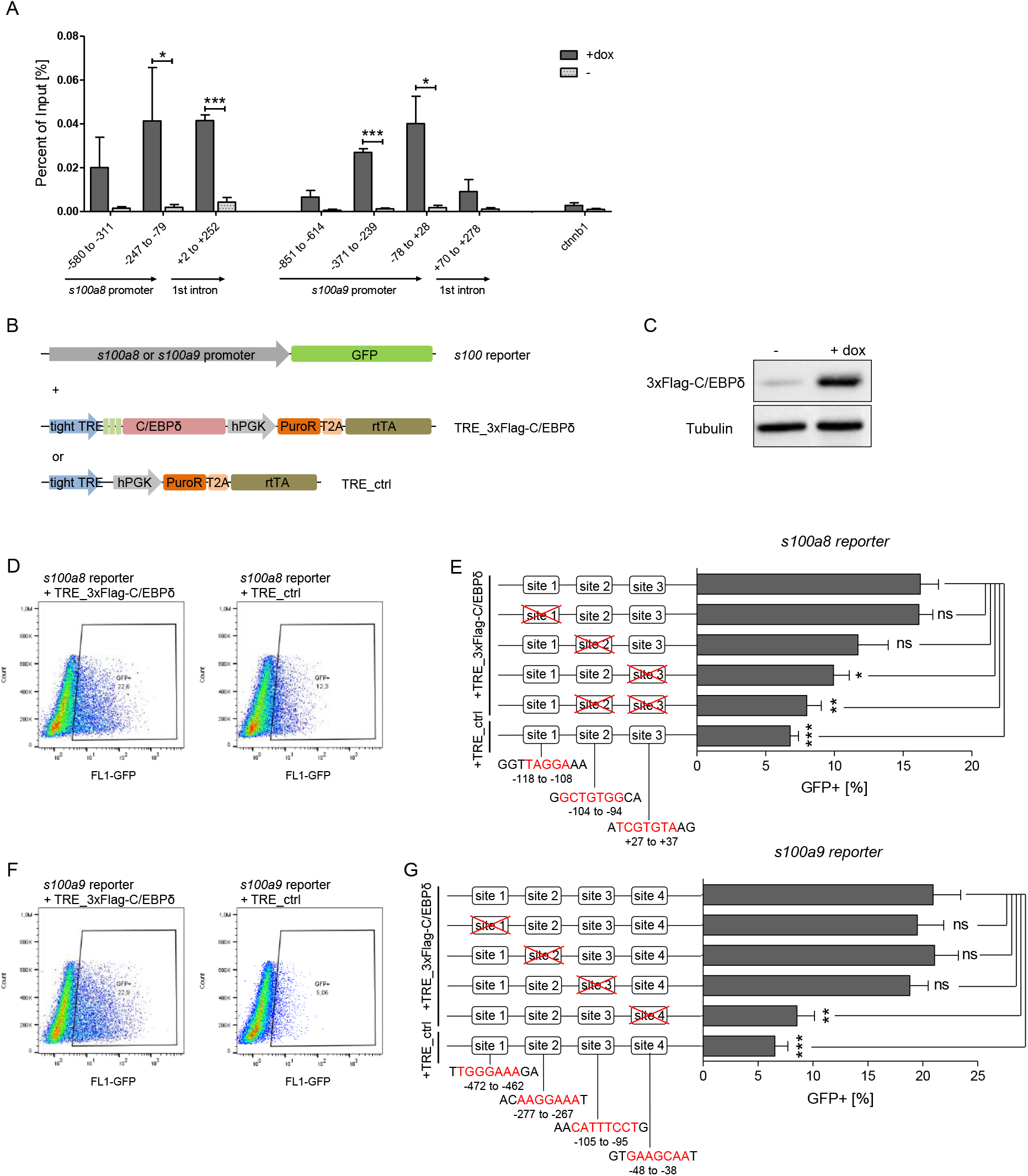
C/EBPδ binds to regions within the *s100a8* and *s100a9* promoters. (A) Chromatin immunoprecipitation was performed in untreated (-) and dox-treated (+dox) TRE_3xFlag-C/EBPδ monocytes on differentiation day 1 using a Flag-antibody. Purified DNA was analysed using primer pairs flanking different s100a8 and s100a9 promoter and intronic regions and a negative control primer pair flanking a random genomic region (n = 3). (B) Co-transfection of vectors carrying constructs for GFP under the *s100a8* or *s100a9* promoter (reporter), together with the doxycycline-dependent 3xFlag-C/EBPδ expression cassette (TRE_3xFlag-C/EBPδ) or a corresponding control vector lacking the 3xFlag-C/EBPδ expression cassette (TRE_ctrl) in HEK293T cells, was performed. (C) Induction of 3xFlag-C/EBPδ upon doxycycline treatment (2 µg/mL, 24h) was analysed by western blot. Representative dot plots from flow cytometry analysis show GFP+ gates of co-transfected HEK293T cells, either using TRE_3xFlag-C/EBPδ or TRE_ctrl together with *s100a8* reporter (D) and with *s100a9* reporter (F) upon doxycycline treatment (2 µg/mL, 24h). Co-transfection of TRE_3xFlag-C/EBPδ and *s100a8* (E) and *s100a9* (G)-reporter plasmids carrying different mutated possible binding sites was performed, analysed 24h post-transfection and compared to co-transfection of TRE_ctrl and s100-reporter plasmid activities. Suggested C/EBP binding sites targeted by depletion are indicated by nucleic acids marked in red (n = 4-5). Values are the means ± SEM. *P < 0.05, **P < 0.01, ***P < 0.001, ns = not significant, by two-tailed Student’s t test.

### Epigenetic landscape on s100 promoter regions reflects S100A8 and S100A9 expression in monocytes

Regulation of gene expression relies on variable factors; among these are chromatin structure and epigenetic features. To measure changes in chromatin accessibility in monocytic-progenitors and in S100A8/A9 expressing monocytes, we performed ATAC-seq of precursor and differentiated ER-Hoxb8 cells. This revealed over 20,000 regions with differential peaks (Figure 6, A). Openness of chromatin between precursor and differentiated cells was highly different, as shown by principal component analysis (Figure 6, B). Among the regions with significantly higher ATAC-seq reads in differentiated samples were the *s100a8* and *s100a9* promoter and enhancer locations (Figure 6, C). Consistent with the changes in chromatin accessibility at *s100* promoter regions during differentiation, we also found changes in histone marks by ChIP. H3K27 acetylation (H3K27ac), a marker for active transcription, was increased at differentiation day 3 in monocytes over precursor cells at *s100a8* (Figure 6, D) and *s100a9* loci (Figure 6, E) in both, WT and C/EBPδ KO cells. In contrast, tri-methylated H3K27 (H3K27me_3_), associated with gene silencing, was overrepresented in precursor cells over differentiated cells at the same loci in WT cells, whereas H3K27me3 marks did not decrease over the course of differentiation in C/EBPδ KO cells. Accordingly, tri-methylated H3K27 was increased in C/EBPδ KO monocytes, compared to the WT counterparts (Figure 6, F and G).

**Figure 6:**
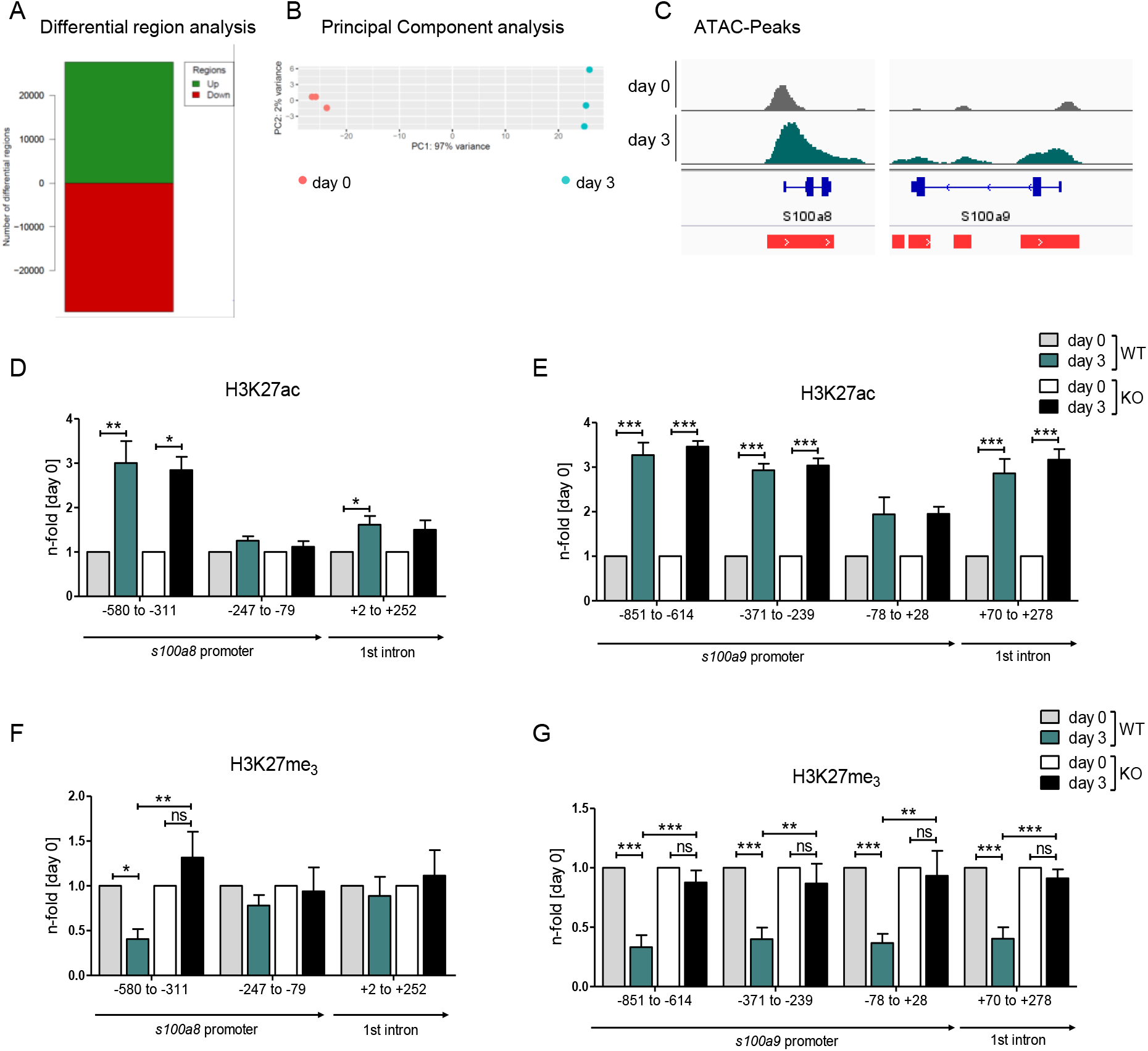
Analysis of chromatin accessibility and epigenetic features within *s100a8* and *s100a9* promoter regions. ATAC sequencing (ATAC-seq) was executed in precursor (day 0) and differentiated (day 3) WT ER-Hoxb8 monocytes. (A) Differential region analysis and (B) principal component analysis (PCA) were performed (n = 3). (C) Representative gene tracks showing ATAC-seq reads of precursor (day 0) and differentiated (day 3) cells at the *s100a8* and *s100a9* gene regions. Red bars beneath genomic locations mark regions with significantly increased ATAC-signals in day 3 samples compared to day 0 samples. Chromatin-Immunoprecipitation was performed using anti-H3K27ac (D, E), anti-H3K27me3 (F, G) in chromatin of precursor (day 0) and differentiated (day 3) WT and C/EBPδ KO (KO) ER-Hoxb8 monocytes. Purified DNA was analysed using primer pairs flanking different *s100a8* (D, F) and *s100a9* (E, G) promoter regions (n = 3-6). N-folds are based on percent of input-values of respective day 0 ChIP-PCR samples. Values are the means ± SEM. *P < 0.05, **P < 0.01, ***P < 0.001, ns = not significant, by one-way ANOVA with Bonferroni correction.

### The histone demethylase JMJD3 drives s100a8/a9 expression in dependency of C/EBPδ

Decreased S100 expression in C/EBPδ-deficient day 3 monocytes is mirrored in the epigenetic landscape by only slightly decreased H3K27ac level, but highly increased H3K27me_3_ level at s100 promoter regions. Erasure of tri-methylation and di-methylation at H3K27 is known to be catalyzed by the histone demethylase JMJD3 (JmjC Domain-Containing Protein 3) (Xiang *et al.*, 2007). We found decreased expression of *jmjd3* in differentiated C/EBPδ KO monocytes, compared to WT cells at the same stage (Figure 7, A). Further, we used the potent JMJD3 inhibitor GSK-J4 (Kruidenier *et al.*, 2012) to block H3K27 demethylation in differentiating ER-Hobx8 cells, and discovered significantly decreased *s100a8* and *s100a9* expression in GSK-J4 treated WT cells (Figure 7, B). These mRNA quantities were comparable to untreated C/EBPδ-deficient monocytes, whereas the effects on *s100a8/a9* expression in GSK-J4-treated C/EBPδ-deficient monocytes, compared to the untreated counterparts, were negligible (Figure 7, B). These effects of GSK-J4 on *s100a8* and *s100a9* expression are in line with increased H3K27me_3_ marks in GSK-J4-treated WT monocytes, compared to untreated WT cells on both, *s100a8* (Figure 7, C) and *s100a9* (Figure 7, D) promoter regions.

**Figure 7:**
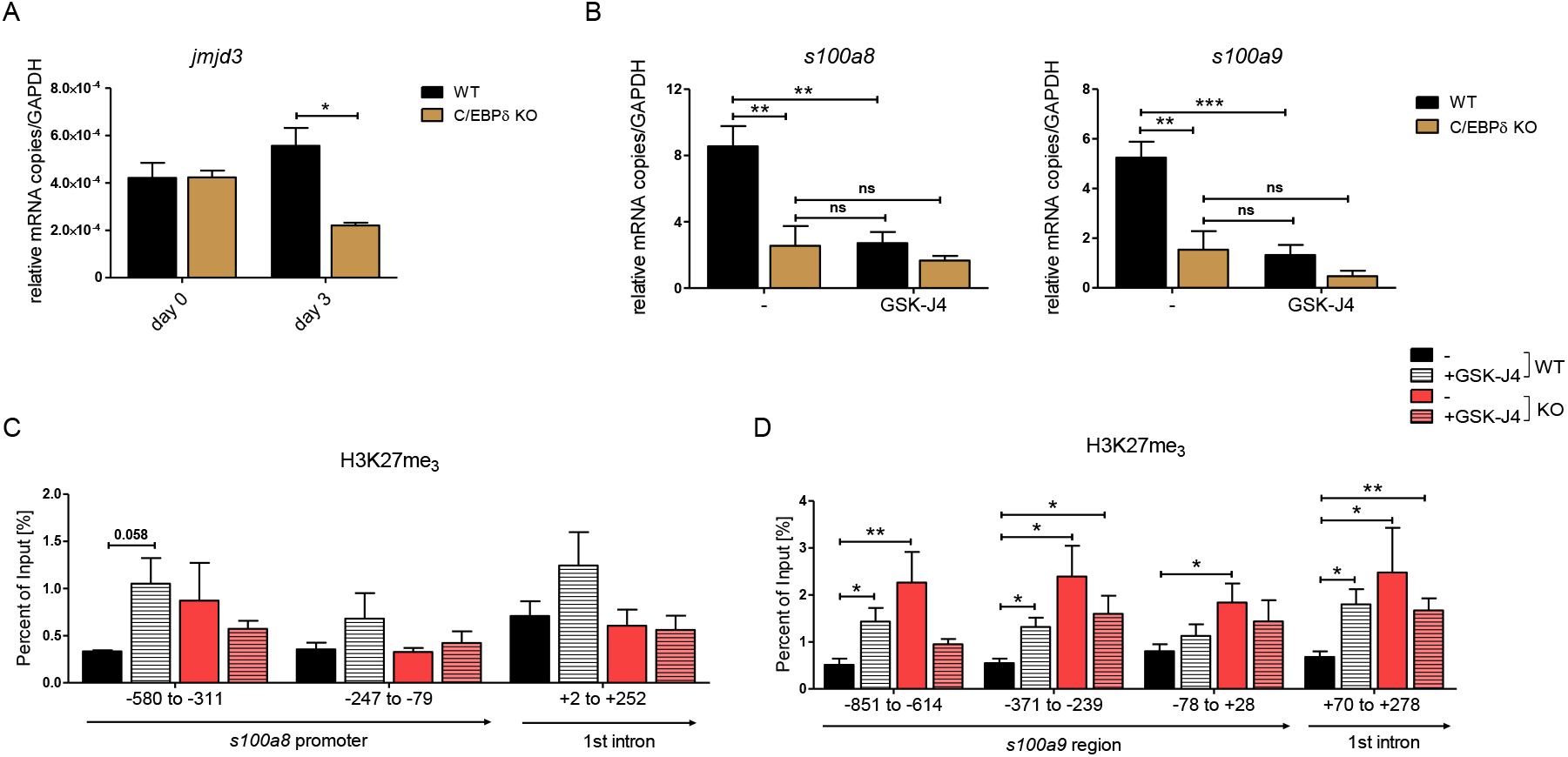
JMJD3-mediated demethylation of H3K27me3 is crucial for *s100a8* and *s100a9* expression. (A) *Jmjd3* mRNA levels of precursor and differentiated WT and C/EBPδ KO ER-Hoxb8 cells were analysed using qRT-PCR (n = 3). (B) WT and C/EBPδ KO ER-Hoxb8 cells were treated with 5 µM GSK-J4 for 3 days during differentiation and *s100a8* and *s100a9* mRNA levels were analysed using qRT-PCR (n = 5). Chromatin-Immunoprecipitation was performed using anti-H3K27me3 and appropriate IgG control antibodies in chromatin of vehicle controls (-) and treated (+GSK-J4) WT and C/EBPδ KO (KO) ER-Hoxb8 monocytes. Purified DNA was analysed using primer pairs flanking different *s100a8* (C) and *s100a9* (D) promoter regions (n = 3-5). Values are the means ± SEM. *P < 0.05, **P < 0.01, ***P < 0.001, ns = not significant, by one-way ANOVA with Bonferroni correction (A, B) or by two-tailed Student’s t test in comparison to WT (-) (C, D).

## DISCUSSION

The ER-Hoxb8 cell system serves as a substitute for murine primary cells of myeloid origin that can be differentiated into phagocytes, such as monocytes and neutrophils. Using this system allows comparison with *in vitro* differentiated primary cells (G. G. Wang *et al.*, 2006) and, therefore, provides an experimental cell model for analysis of S100A8 and S100A9 expression. Although the alarmin is regarded as key factors in various inflammatory conditions (Foell and Roth, 2004), cancer types (Cross *et al.*, 2005) and cardiovascular diseases (Frangogiannis, 2019), little is known about their transcriptional regulation. The serum concentrations of alarmins correlate with disease severity and activity and, hence, they are reliable biomarkers for monitoring several inflammatory diseases(Foell *et al.*, 2004; Ehrchen *et al.*, 2009). The expression levels of S100A8 and S100A9 differ extremely during myeloid differentiation and the promotors of their genes represent probably one of the most dynamic regulatory elements in the myeloid lineage. Whereas both proteins are completely absent in myeloid precursor cells, they are highly expressed in monocytes and neutrophils, which suggests that highly dynamic regulatory mechanisms drive S100A8 and S100A9 expression.

The CRISPR/Cas9-mediated KO screening approach based on a lentiviral pooled library has been used so far to investigate various mechanisms, such as immunity-related pathways and cancer-modulating events (Kweon and Kim, 2018). In this study, our unbiased genome-wide screening approach allowed the identification of C/EBPδ as a factor involved in S100A9 regulation during murine monocyte differentiation. We further focused our investigations on *cebpd* because this gene was in the top list of the robust rank aggregation (RRA) scores and showed the highest numbers of guide RNAs with efficient effects on S100A9 expression in our screening. This redundancy of independent parameters helped to distinguish true positive from false positive hits. Furthermore, a robust phenotype-of-interest, such as a clear S100A9 protein signal at day 3 of monocyte differentiation, allowed reliable negative selection in Cas9-library monocytes. Selection of remaining cells served as a reference control to distinguish true from false positives. The specificity of our selection procedure was confirmed at the protein level by western blot analysis of sorted cell populations. CRISPR/Cas9-based functional genomic screening has been shown to be highly specific, thereby causing fewer cases of false positives in direct comparison with knockdown analysis by RNA interference (Shalem *et al.*, 2014). We were now able to identify a novel regulator of S100A8 and S100A9 using this unbiased method. By pre-gating on CD11b^+^Ly-6C^+^ monocytes we revealed C/EBPδ as a specific and differentiation-independent regulator of S100A8 and S100A9, excluding pathways linked to general functions or development of phagocytes.

We confirmed that the transcription factor C/EBPδ is a direct regulator of S100A8 and S100A9 in murine monocytes in independent approaches. C/EBPδ and S100A8/A9 are co-expressed in differentiating monocytes, and induction of C/EBPδ clearly showed that the expression of S100A8 and S100A9 was up-regulated by the presence of C/EBPδ. This evidence was further supported by increased S100A8/A9 levels caused by deletion of ATF3 and FBXW7, which are natural inhibitors of C/EBPδ.

The specificity of our approach was further confirmed by the fact that deficiency of several transcription factors, such as STAT3, KLF5, IRF7 and C/EBPβ, described as S100A8/A9 regulators in previous studies (Kuruto-Niwa *et al.*, 1998; Fujiu, Manabe and Nagai, 2011; Lee *et al.*, 2012; Liu *et al.*, 2016; Yang *et al.*, 2017), did not affect S100A8 and S100A9 expression in our ER-Hoxb8 monocytes.

Our ChIP data clearly showed that C/EBPδ specifically binds within *s100a8* and *s100a9* promoter regions. Co-transfection of an inducible C/EBPδ-construct and *s100a8* and *s100a9* reporter constructs not only demonstrated *s100* promoter activation due to C/EBPδ expression, but also revealed functional relevance of specific binding sites, via promoter bashing, that are located exactly within the stated promoter regions. The two DNA-motifs for specific C/EBPδ responses on the *s100a8* promoter regions did not share the core sequence 5’-C/G GCAAT-3’ that we found within the *s100a9* promoter region in our study. The latter has been described in three other promoters, the human *pparg2* promoter (Lai *et al.*, 2008), the murine and human *cebpd* promoter itself (Wang *et al.*, 2021) and the human *cox-2* promoter (J. M. Wang *et al.*, 2006). We were able to show that the functionally relevant C/EBPδ binding sites within the S100 promoters lie within genome regions which switch from closed to open chromatin states during monocyte differentiation, and concomitant induction of S100 expression as examined by ATAC-seq.

Our chromatin accessibility data on *s100a8* and *s100a9* promoter regions reflected active *s100a8 and s100a9* transcription and was supported by the characterization of the epigenetic landscape using H3K27ac and H3K27me_3_ marks. The fact that H3K27me_3_ marks were strongly decreased in WT monocytes, but not in precursors or C/EBPδ-deficient monocytes, showed the indispensability of H3K27 demethylation for S100A8/A9 expression. Moreover, our data demonstrated that the Jumonji C family histone demethylase JMJD3 regulates S100A8/A9 expression by erasure of H3K27me_3_ in dependency of C/EBPδ, which was confirmed by GSK-J4-mediated inhibition of JMJD3 activities. Neither a link of C/EBPδ, nor of S100A8/A9 and JMJD3 has been published yet. It has been shown that histone demethylase activities of recombinant JMJD3 on mono-nucleosome substrates is relatively low in contrast to higher activities on bulk histones (Lan *et al.*, 2007), suggesting that further factors, such as C/EBPδ, are involved in chromatin binding. Several studies highlight JMJD3 as a regulator of innate immune responses, especially via NF-κB-mediated inflammation in macrophages (Na *et al.*, 2016, 2017; Davis *et al.*, 2020). Accordingly, knockdown of JMJD3 affected mainly inflammatory response networks in monocytic THP-1 cells (Das *et al.*, 2012) and blocked activation of the NLRP3 inflammasome in bone marrow-derived macrophages (Huang *et al.*, 2020). GSK-J4 treatment of mice attenuated disease progression and inflammatory activities in several mouse models for inflammatory diseases, such as arthritis (Jia *et al.*, 2018), colitis (Huang *et al.*, 2020) and EAE (experimental autoimmune encephalomyelitis) (Doñas *et al.*, 2016). Accordingly, GSK-J4 treatment of our ER-Hoxb8 monocytes reduced expression of the proinflammatory alarmin S100A8/A9, which have been shown to drive the inflammatory process of arthritis (Van Lent *et al.*, 2012). With our study, we have taken a step forward to uncover the role of epigenetic features on S100A8 and S100A9 expression and, thereby, on inflammatory conditions in murine monocytes.

We were also able to demonstrate an association of C/EBPδ and S100A8/A9 expression in the context of human cardiovascular disease. We did not only show that expression of these molecules show a significant positive correlation to each other but also to the manifestation of stable coronary artery disease (sCAD) and myocardial infarction (MI) in patient derived PBMCs. Moreover, expression of C/EBPδ and S100A8/A9 showed an even stronger association with classical, proinflammatory monocytes (CD14^++^CD16^-^), compared to non-classical (CD14^+^CD16^++^) and intermediate (CD14^++^CD16^+^) monocytes. The endogenous antagonists of C/EBPδ, ATF3 and especially FBXW7, showed a negative correlation of their expression pattern in these monocyte subpopulations. Interestingly, inflammatory monocytes with phagocytic and proteolytic activities have been reported to show an early peak at infarct sites, which are followed by infiltration of non-classical, anti-inflammatory monocytes (Nahrendorf *et al.*, 2007; Dutta and Nahrendorf, 2015). Genetic deletion of S100A8/A9 was reported to attenuate MI and improve cardiac function in murine models. In contrast, overexpression of S100A9 in mice increased infarct size and mortality, and treatment with recombinant S100 proteins raised influx of immune cells into the infarct area (Li *et al.*, 2019; Sreejit *et al.*, 2020). Moreover, serum concentrations of S100A8/A9 are known to be highly sensitive and prognostic markers for myocardial injury (Aydin *et al.*, 2019). Taken together, these data indicate that the C/EBPδ-S100-alarmin axis drives a clinically relevant pathomechanism in cardiovascular disease and probably other inflammatory driven conditions.

There are several published reports suggesting a biomedical relevance of the link between the C/EBPδ and S100A8/A9 alarmin under other inflammatory conditions as well. For example, C/EBPδ has been shown to play a role in the pathogenesis of psoriasis (Lan *et al.*, 2020) and in acute inflammatory signaling by regulating COX-2 (Wadleigh *et al.*, 2000), IL-6 (Litvak *et al.*, 2009) and TLR4 (Balamurugan *et al.*, 2013). Analysis of the genome-wide transcription pattern of monocytes revealed IL-6 as the top gene induced by S100 alarmin stimulation via interaction with TLR4 (Fassl *et al.*, 2015), and targeted deletion of S100A9 ameliorated inflammation in a murine psoriasis model(Zenz *et al.*, 2007). Additionally, C/EBPδ levels were elevated in mouse models and patients of Alzheimer’s disease (AD) (Li *et al.*, 2004; Ko *et al.*, 2012) and rheumatoid arthritis (RA) (Nishioka *et al.*, 2000; Chang *et al.*, 2012). In mouse models of AD, down-regulation (Ha *et al.*, 2010) and deficiency of S100A9 (Kummer *et al.*, 2012) had therapeutic effects on disease activity. Also in human studies, S100A9 was found to be associated with AD pathogenesis (Shepherd *et al.*, 2006). Beyond that, S100A8 and S100A9 are known key players in the pathogenesis of arthritis in murine models (Van Lent *et al.*, 2012). Gene expression profiling of blood cells from RA patients receiving anti-TNF-α-based treatment showed that both C/EBPδ and S100A8 were downregulated by the treatment (Meugnier *et al.*, 2011). Uncontrolled activity of S100A8/A9 alarmins drives TNF-induced arthritis in mice (Vogl *et al.*, 2018).

In the context of human RA, the expression and serum concentrations of S100A8/A9 correlate very well with disease activity and are the first predictive marker for disease relapses in juvenile patients, and of the responses to therapy in juvenile and adult patients (Moncrieffe *et al.*, 2013; Choi *et al.*, 2015). However, no direct molecular or functional link between S100A8/A9 and C/EBPδ in arthritis has yet been reported.

## MATERIAL & METHODS

### Cell culture

ER-Hoxb8 cells were generated as described earlier (G. G. Wang *et al.*, 2006) and grown in RPMI medium (Thermo Fisher Scientific) supplemented with 10% FBS (Biowest), 1% penicillin/streptomycin solution (Sigma-Aldrich), 1% glutamine solution (Thermo Fisher Scientific) 40 ng/ml rmGM-CSF (ImmunoTools) and 1 µM β-estradiol (Sigma-Aldrich). For differentiation, precursor cells were washed and incubated in estradiol-free medium containing 40 ng/ml rmGM-CSF for several days. HEK293T were grown in DMEM (Thermo Fisher Scientific) supplemented with 10% FBS (Biowest) and 1% penicillin/streptomycin solution (Sigma-Aldrich), 1% glutamine solution (Thermo Fisher Scientific), and 1% sodium pyruvate (Merck). All cell lines were cultured at 37 °C, 5% CO_2_ and routinely screened and found negative for mycoplasma contamination in a PCR-based assay (PromoCell).

### Cell generation and manipulation

WT, C/EBPδ KO (kindly provided by Esta Sterneck, National Cancer Institute, Frederick, MD, USA) (Sterneck *et al.*, 1998) and Cas9 expressing (Chiou *et al.*, 2015) cells originated from corresponding mice. FBXW7, ATF3, STAT3, KLF5, IRF7 and C/EBPβ KO ER-Hoxb8 cells were generated using CRISPR/Cas9 as described earlier (Gran *et al.*, 2018). The oligos for gRNA cloning are listed in Supplementary Table S2. For lentiviral production, the lentiGuide-Puro (for GeCKO screen), lentiCRISPRv2-gRNA (for single KO cell lines) or TRE_3xFlag-C/EBPδ was co-transfected into HEK293T cells, together with the packaging plasmids pCMV-VSV-G (AddGene, #8454) and psPAX2 (AddGene, #12260). For transduction of ER-Hoxb8 cells, cells were incubated with lentiviral particles and 8 µg/ml polybrene (Sigma-Aldrich) for 1 hour upon spinfection and selected for several days using puromycin (InvivoGen). For transfection of HEK293T cells, the cells were seeded one day prior to transfection. Then, cells were co-transfected with TRE_3xFlag-C/EBPδ and *s100a8* reporter or *s100a9* reporter using the Lipofectamine™ 3000 Transfection Reagent (Thermo Scientific) according to the manufacturer manual. For inhibition of JMJD3-activities, cells were treated using 5 µM GSK-J4 HCl (SellekChem) for 3 days. To induce *cebpd* in TRE_3xFlag-C/EBPδ ER-Hoxb8 cells or transfected HEK293T cells, cells were treated using 2 µg/mL doxycycline (Sigma-Aldrich) for 24 hours.

### GeCKO-library screening

Amplification of mouse CRISPR Knockout pooled library (GeCKO v2) in lentiGuide-Puro plasmid, purchased from AddGene (#1000000053) (Sanjana, Shalem and Zhang, 2014), was performed as described (Joung *et al.*, 2017). Cas9 expressing ER-Hoxb8 cells, transduced with library lentiviral particles at a MOI of 0.4, were differentiated to day 3. Intracellular S100A9 was stained with a S100A9-FITC coupled antibody using the Foxp3/Transcription Factor Staining Buffer Set (eBioscience). Cells with no/lower S100A9 expression (hits) and cells with normal S100A9 expression (reference) were sorted using a SH800S Cell Sorter (Sony, Minato, Japan) and DNA was purified by phenol-chloroform extraction. Next generation sequencing was performed as described earlier (Joung *et al.*, 2017). Briefly, sgRNA library for next generation sequencing was prepared via PCR using primers amplifying the target region with Illumina adapter sequences (Supplementary Table S3), the purified DNA and the NEBNext® High-Fidelity 2X PCR Master Mix (NEB). PCR reactions were pooled and purified using the QIAquick PCR Purification Kit (Qiagen). Size and quantity was determined using the Bioanalyzer High Sensitivity DNA Analysis Agilent High Sensitivity DNA Kit (Agilent). Samples were sequenced according to the Illumina user manual with 80 cycles of read 1 (forward) using the NextSeq 500/550 High Output Kit v2.5 (75 Cycles) (Illumina) with the 20% PhiX spike in Illumina PhiX control kit (Illumina).

### Cloning and plasmid production

#### TRE_3xFlag-C/EBPδ and TRE_ctrl

The pcDNA 3.1 (-) mouse C/EBPδ expression vector (AddGene, #12559) and annealed oligonucleotides (Supplementary Table S4) were digested using *Xba*I and *EcoR*I and then ligated. Using primers carrying restriction enzyme recognition sites (Supplementary Table S4), the 3xFlag-C/EBPδ expression cassette was amplified. The resulting amplicon and the pCW57.1 mDux-CA target vector (AddGene, #99284) (Whiddon *et al.*, 2017) were digested using *Nhe*I and *Age*I and subsequently ligated. TRE_ctrl was produced by digesting TRE_3xFlag-C/EBPδ using *Nhe*I and *Age*I and by subsequent blunting of ends by 3’ overhang removal and fill-in of 3’ recessed (5’ overhang) ends using DNA Polymerase I, Large (Klenow) Fragment (NEB) prior to ligation.

#### S100a8 and s100a9 reporter

To construct *s100-*reporter vectors, 1500 bp upstream of *s100a8* and 1800 bp upstream of *s100a9* transcription start sites were amplified from genomic mouse DNA. Using primers carrying restriction enzyme recognition sites (Supplementary Table S5), promoter regions were amplified and cloned into pLenti CMV GFP Blast vector (AddGene, #17445) (Campeau *et al.*, 2009) using *Xba*I and *Cla*I. Resulting *s100*prom-GFP constructs were cloned into MSCV-PIG-empty vector (AddGene, #105594) (Xu *et al.*, 2018) by digestion with *Nsi*I and *Cla*I together with the MSCV-backbone to exchange IRES-GFP-cassette with *s100a8/a9*prom-GFP-cassette and subsequent ligation. Proposed C/EBP DNA binding sites within *s100a8* and *s100a9* promoter regions were identified using the AliBaba2.1 net-based transcription factor binding site (TFBS) search tool (Grabe, 2002), and were mutated by deleting 6-7 base pairs using the QuikChange II XL Site-Directed Mutagenesis Kit (Agilent Technologies). The primers used for mutagenesis are listed in Supplementary Table S6. Plasmids were produced in DH5α cells and purified using the PureLinkTM HiPure Plasmid Midiprep Kit (ThermoScientific).

### Quantitative reverse transcription polymerase chain reaction (qRT-PCR)

RNA was isolated using a NucleoSpin Extract II Isolation Kit (Macherey Nagel). The mRNA expression of selected genes was measured by qRT-PCR as described earlier (Heming *et al.*, 2018). The primers used are listed in Supplementary Table S7. The relative expression level of each target gene was analysed using the 2−ΔΔCq method and was normalised to GAPDH.

### Chromatin immunoprecipitation (ChIP)

For chromatin preparation, progenitor and differentiated ER-Hoxb8 cells were fixed using 1% formaldehyde for 5 minutes and reaction was stopped by adding 125 mM glycine. Chromatin was extracted as previously described (Fujita and Fujii, 2013). Approximately 1-5% of chromatin served as the input sample. DNA from input samples was isolated using phenol-chloroform extraction as described earlier (Heming *et al.*, 2018). For immunoprecipitation, 3 µg antibody against Flag (Sigma-Aldrich, #F1804), H3K27ac (Abcam, #ab4729), H3K27me_3_ (Cell Signaling Technology, #9733), normal Rabbit IgG (Cell Signaling, #2729) or Mouse IgG1, κ Isotype control (Biolegend, #400102) was conjugated to 900 µg magnetic Dynabeads-Protein G (Thermo Scientific, Waltham, MA) at 4 °C overnight. Sonicated chromatin was added to AB-conjugated Dynabeads and incubated at 4 °C overnight. The Dynabeads were washed as described earlier (Fujita and Fujii, 2013). For elution, Dynabeads were incubated twice with elution buffer (0.05 M NaHCO_3_, 1% SDS) at 65 °C for 15 minutes. DNA from eluates was isolated using phenol-chloroform extraction as with input samples. Values were taken into account only when the amount of DNA pulled down by using the antibody of interest was more than 5-fold increased over DNA pulled down by using IgG antibodies. The primers used for ChIP-PCR are listed in Supplementary Table S8.

### Assay for Transposase-Accessible Chromatin using sequencing (ATAC-seq)

Precursor and day 3 differentiated WT ER-Hoxb8 cells were harvested, washed and cryopreserved in 50% FBS/ 40% growth media/ 10% DMSO using a freezing container at −80 °C overnight. Cells were shipped to Active Motif to perform ATAC-seq as previously described (Buenrostro *et al.*, 2013).

### Measurements of S100A8/A9 protein level

The S100A8/A9 protein concentrations were measured using an in-house S100A8/A9 enzyme-linked immunosorbent assay (ELISA), as previously described (Vogl *et al.*, 2014).

### Immunoblotting

Cells were lysed in M-PER™ Mammalian Protein Extraction Reagent (Thermo Scientific, Waltham) containing a protease inhibitor mixture (Sigma-Aldrich). Protein concentration was determined, and equal amounts (15-30 µg) were run on a SDS-PAGE. After blotting on a nitrocellulose membrane, the membrane was incubated overnight with primary antibodies against: polyclonal rabbit S100A8 and S100A9 antibodies (originated from our own production (Vogl *et al.*, 2014)), GAPDH (Cell Signaling Technology), α/β-Tubulin (Cell Signaling Technology) and Flag (Sigma-Aldrich). Membranes were incubated with a HRP-linked secondary antibodiy (Agilent, Santa Clara, CA) for 1 hour. Chemiluminescence signal was detected using ChemiDoc XRS+ (Bio-Rad) together with ImageJ (National Institutes of Health) to quantify the signal intensity.

### Phagocytosis of Latex Beads

FluoSpheres polystrene microspheres (ThermoScientific) were shortly sonicated in a bath sonicator and added to cells at a ratio 1:10 for 2 h at 37 °C. The rate of phagocytosis was determined by flow cytometry using Navios (Beckmann Coulter).

### Oxidative Burst

Cells were stimulated using 10 nM PMA (Abcam) for 15 minutes or left untreated. After incubation, 15 μM DHR123 (Sigma-Aldrich) were added for another 15 min. The fluorescence signal was analysed using flow cytometry (Navios, Beckmann Coulter).

### RNA-sequencing (RNA-seq)

#### Study population

For this study, we used bulk mRNA-sequencing (RNA-seq) data of peripheral blood mononuclear cells (PBMCs) and monocytes from two subsets of participants in the German BioNRW Study. BioNRW actively recruits patients undergoing coronary angiography for the diagnosis and percutaneous coronary intervention of coronary artery disease, as well as age and gender matched healthy control individuals without history of cardiovascular disease, all aged 18-70 years old. Patients receive standard cardiovascular care and medication (ACE-inhibitor, AT1-receptor blocker, β-blocker, diuretics, statin), according to current guidelines. Here, we included a total of 42 patients with stable coronary artery disease (sCAD) or acute myocardial infarction (MI), as well as 39 of the corresponding age and sex matched controls. The BioNRW Study is conducted in accordance with the guidelines of the Declaration of Helsinki. The research protocol, including the case report forms, was approved by the local ethics committee (#245-12). Written informed consent was obtained from all study participants.

#### Blood collection and isolation of PBMCs

In case of MI, blood samples were collected during the first 4 days following the event. EDTA blood was drawn from each subject by venipuncture. Sample processing followed within two hours. PBMCs were obtained from 40 mL blood by density gradient centrifugation (Ficoll; Biochrom). Lymphocytes were collected and washed twice with PBS. The pellet was re-suspended in freezing medium Cryo-SFM (PromoCell) and cryopreserved.

#### Isolation of monocyte subpopulations

After washing, PBMCs were stained with anti-human antibodies specific for CD2 (PE, RPA-2.10, T-cell marker), CD14 (APC, M5E2, monocyte subset differentiation), CD15 (PE, HIM1, granulocyte marker), CD16 (PE-Cy7, 3G8, monocyte subset differentiation), CD19 (PE, HIB19, B-cell marker), CD56 (PE, MY31, NK-cell marker), CD335 (PE. 9E2, NK-cell marker), HLA-DR (FITC, TU36, antigen-presenting cells) (all from BD Biosciences), as reported by Cros et al. (2010). Cells were acquired on a FACS LSR II flow cytometer (BD Biosciences) and analysed using FlowJo software version 10 (Treestar Inc.). For sorting of monocyte subsets, PBMCs were stained and sorted on a MoFlo Astrios cell sorter (Beckman Coulter). Cells were sorted in 1 mL of Isol-RNA lysis reagent (5-Prime GmbH) and frozen at –80 °C. To avoid gender-specific effects, 3 representative male samples of each BioNRW diagnostic group (sCAD, MI and control) were selected to be subjected to cell sorting and subsequent RNA isolation.

#### Differential expression analysis in PBMCs and monocytes

For mRNA profiling of PBMCs and monocyte subpopulations using RNA-Seq, mRNA was enriched using the NEBNext® Poly(A) Magnetic Isolation Module (NEB), followed by cDNA NGS library preparation (NEBNext® Ultra RNA Library Prep Kit for Illumina, NEB). The size of the resulting libraries was controlled by the use of a Bioanalyzer High Sensitivity DNA Kit (Agilent Technologies) and quantified using the KAPA Library Quantification Kit for Illumina (Roche). Equimolar, appropriately pooled libraries were sequenced in a single read mode (75 cycles) on a NextSeq500 System (Illumina) using v2 chemistry, yielding in an average QScore distribution of 92% ≥ Q30 score. They were subsequently demultiplexed and converted to FASTQ files using bcl2fastq v2.20 Conversion software (Illumina). Data was quality controlled using FASTQC software and trimmed for adapter sequences using Trimmomatic (Bolger, Lohse and Usadel, 2014).

### General statistics

The statistical significance of the data was determined using Prism 5.0 software (GraphPad Software, CA, USA). Analyses between two groups were performed using an unpaired two-tailed Student’s t-test. Comparisons among three or more groups were performed by using one-way ANOVA, followed by Bonferroni’s multiple means tests for comparing all pairs of columns. Differences were considered statistically significant at a probability (p-value) of <0.05.

### Bioinformatics analysis

#### GeCKO-library screening

Analysis of counting the reads for each gRNA and differential analysis was performed using Model-based Analysis of Genome-wide CRISPR-Cas9 Knockout (MaGeCK) 0.5.9.3, a computational tool to identify important genes from GeCKO-based screens (Li *et al.*, 2014). A modified RRA (Robust Rank Aggregation) method with a redefined *ρ* value was used. Former RRA computed a significant P-value for genes in the middle of gRNA ranked list and thereby introducing false positives because the assumption of uniformity is not necessarily satisfied in real applications. Thus, top ranked % gRNAs were selected if their negative binomial P-values were smaller than a threshold, such as 0.05. If j of the n gRNAs targeting a gene were selected, then the modified value is defined as *ρ* = min (p1, p2,…,pj), where j ≤ n. This modified RRA method could efficiently remove the effect of insignificant gRNAs in the assessment of gene significance. A permutation test where the gRNAs were randomly assigned to genes was performed to compute a P-value based on the *ρ* values. By default, 100 x n g permutations are performed, where n g is the number of genes. We then compute the FDR from the empirical permutation P-values using the Benjamini-Hochberg procedure.

#### ATAC-seq

Sequence analysis was performed by mapping the paired-end 42 bp sequencing reads (PE42) generated by Illumina sequencing (using NextSeq 500) to the genome using the BWA algorithm with default settings (“bwa mem”). Only reads that passed Illumina’s purity filter, aligned with no more than 2 mismatches, and mapped uniquely to the genome were used in the subsequent analysis. In addition, duplicate reads (“PCR duplicates”) were removed. For Peak finding, genomic regions with high levels of transposition/tagging events were determined using the MACS2 peak calling algorithm (Zhang *et al.*, 2008). Both reads (tags) from paired-end sequencing represent transposition events, both reads were used for peak-calling but treated a single, independent reads. Fragment density was determined by dividing the genome into 32 bp bins and by defining number of fragments in each bin. For this purpose, reads were extended to 200 bp, which was close to the average length of the sequenced library inserts. Differential regions were determined with the DESeq2 bioconductor package (Love, Huber and Anders, 2014) with absolute log2FC > 0.3 and an FDR corrected p < 0.1.

#### RNA seq

The resulting reads were mapped to the human reference genome builds hg19 (monocytes) or hg38 (PBMCs) using Tophat2 (Kim *et al.*, 2013) or HISAT2 v2.1.0 (Kim *et al.*, 2019), counted by using the R package GenomicAlignments (Lawrence *et al.*, 2013) or HTSeq v0.11.2 (Anders, Pyl and Huber, 2015), and followed by differential expression analysis using DEseq2 (Love, Huber and Anders, 2014). The PBMCs dataset used for analysis consisted of 72 individuals, from which 36 were sCAD/MI cases and 36 were controls (21 females and 15 males in each group, mean age: 50.8 ± 12.3 years), while the monocytes dataset contained read counts of classical, intermediate and non-classical monocyte subpopulations from 9 male individuals (3 MI, 3 sCAD and 3 controls. One sCAD non-classical monocyte sample had to be excluded from analysis due to low mapping rate; therefore, the monocytes dataset used for analysis contained 26 samples. Genes were considered differentially expressed at adjusted p<0.05 (Benjamini-Hochberg method). R was used to perform Pearson correlation tests and generate plots for the genes of interest from the normalised count data.

## CONCLUSION

We found that the transcription factor C/EBPδ drives expression of the abundant alarmins S100A8 and S100A9, and demonstrated that C/EBPδ binding to specific sites on *s100a8* and *s100a9* promoter regions also induced changes in chromatin accessibility via JMJD3-mediated demethylation of H3K27me_3_ marks, which includes a so far unknown link. Due to the high relevance of S100A8/A9 alarmin expression in many inflammatory diseases, our findings may point to novel molecular targets for innovative anti-inflammatory therapeutic approaches.

## ABBREVIATIONS

ATAC-seq: Assay for Transposase-Accessible Chromatin using sequencing
ATF3: activating transcription factor 3
C/EBPβ/δ: CCAAT/Enhancer-Binding-Protein β/δ
CAD: coronary artery disease
Cas9: CRISPR associated protein 9
cDNA: complementary DNA
ChIP: chromatin immunoprecipitation
COX-2: cyclooxygenase-2
CRISPR: clustered regularly interspaced short palindromic repeats
DAMPs: danger-associated molecular patterns
FBXW7: F-Box And WD Repeat Domain Containing 7
GeCKO: Genome-Scale CRISPR/Cas9 Knockout
GFP: green fluorescent protein
GM-CSF: granulocyte-macrophage colony-stimulating factor
IL-6: Interleukin-6
IRF7: interferon regulatory factor 7
JMJD3: JmjC Domain-Containing Protein 3
KLF5: krüppel-like factor 5
KO: Knockout
LPS: lipopolysaccharide
MaGECK: Model-based Analysis of Genome-wide CRISPR-Cas9 Knockout
MI: myocardial infarction
MSCV: murine stem cell virus
MRP8/14: myeloid-related protein 8/14
NGS: Next-generation Sequencing
qRT-PCR: Quantitative reverse transcription polymerase chain reaction
RA: rheumatoid arthritis
ROS: reactive oxygen species
RRA: robust rank aggregation
STAT3: signal transducer and activator of transcription 3
TLR4: toll-like receptor 4
TNF-α: tumor necrosis factor-α
TRE: Tet Response Element
WT: wildtype

## AUTHORSHIP

Contribution: S.-L.J.-S., A.I.C., M.S., J.R., and O.F. conceived and designed the experiments; S.-L.J.-S., A.W., J.W. and O.F. performed experiments; B.M. and B.S. recruited the patients and provided the PBMCs and monocyte subsets from the Bio.NRW study; S.-L.J.-S., M.H.-R., L.M., A.W., M.S., T.V., J.R. and O.F. analysed the data; and S.-L.J.-S., J. R., and O.F. wrote the manuscript.

## ACKNOWLEDGEMENTS

The authors thank Ursula Nordhues, Heike Berheide, Eva Nattkemper, Heike Harter, Elvira Barg and Marianne Jansen-Rust for their excellent technical support, and Esta Sterneck (Center for Cancer Research, National Cancer Institute, Frederick, MD) for providing the C/EBPδ KO mice.

This work was supported by grants the Interdisciplinary Center of Clinical Research at the University of Münster (Ro2/023/19, Vo2/011/19), the German Research Foundation CRC 1009 B8, B9 and Z2, CRU 342 P3 and P5 and RO 1190/14-1 (to J. Roth and T. Vogl) and by the EU EFRE Bio.NRW programme (005-1007-0006) to M.S. The funders had no role in the study design, data collection and analysis, decision to publish, or preparation of the manuscript.

## RESOURCE AVAILABILTY

### lead contact

Further information and requests for resources and reagents should be directed to and will be fulfilled by the lead contact, Johannes Roth

### materials availability

Any resource and reagent in this paper is available from the lead contact upon request.

### data and code availability

Human RNA-seq data have been deposited at the NCBI’s BioProject Database with the ID 706411.

Jauch-Speer SL, Wolf J, Herrera-Rivero M, Martens L, Imam Chasan A, Witten A, Markus B, Schieffer B, Vogl T, Stoll M, Roth J and Fehler O (2021) **GeCKO screen** ID PRJNA754262 at the NCBI’s Database: https://dataview.ncbi.nlm.nih.gov/object/PRJNA754262?reviewer=6l1pv4hrhhb8mcbvcne6psjulv

Jauch-Speer SL, Wolf J, Herrera-Rivero M, Martens L, Imam Chasan A, Witten A, Markus B, Schieffer B, Vogl T, Stoll M, Roth J and Fehler O (2021) **ATAC-seq in precursor and differentiated ER-Hoxb8 cells** ID PRJNA755208 at the NCBI’s Database: https://dataview.ncbi.nlm.nih.gov/object/PRJNA755208?reviewer=ivcq37bj1esdf9n7vl6l83m3 ed

## SUPPLEMENTAL ITEMS

### FIGURES

**Figure 2 – figure supplement 1:**
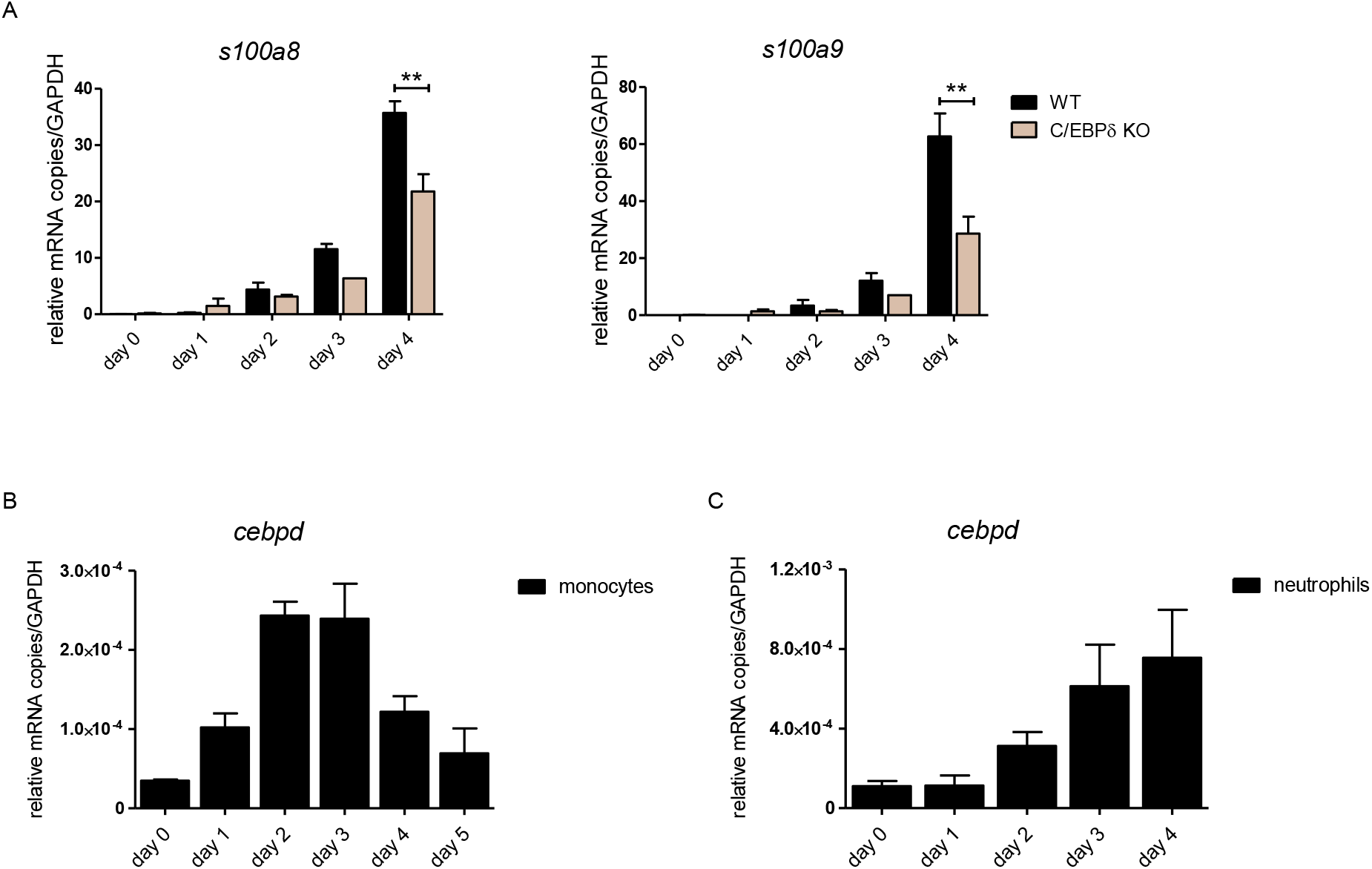
S100A8, S100A9 and C/EBPδ-expression kinetics in ER-Hoxb8 cells.

**Figure 2 – figure supplement 2:**
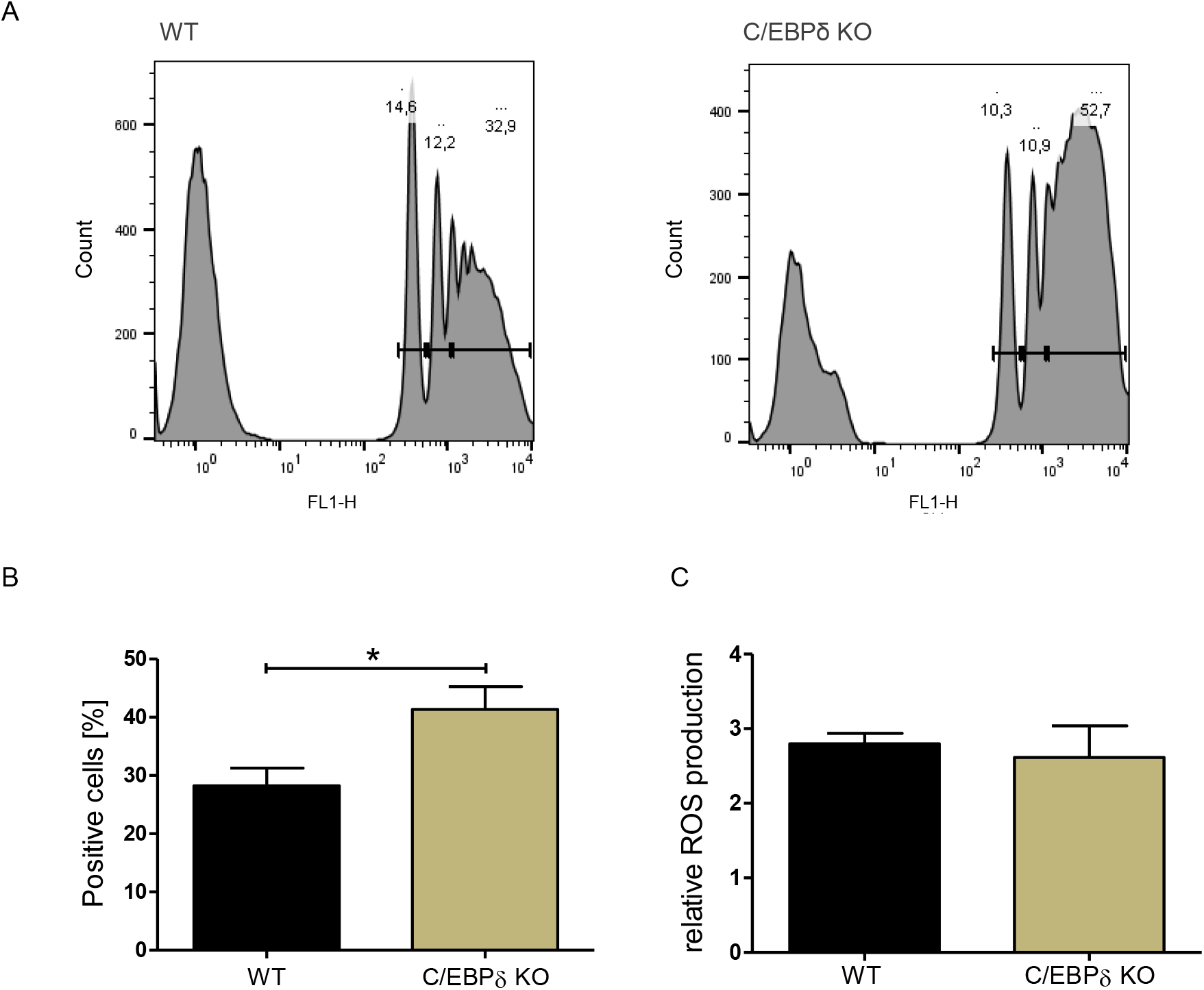
Functional properties of WT and C/EBPδ KO ER-Hoxb8 monocytes.

**Figure 2 – figure supplement 3:**
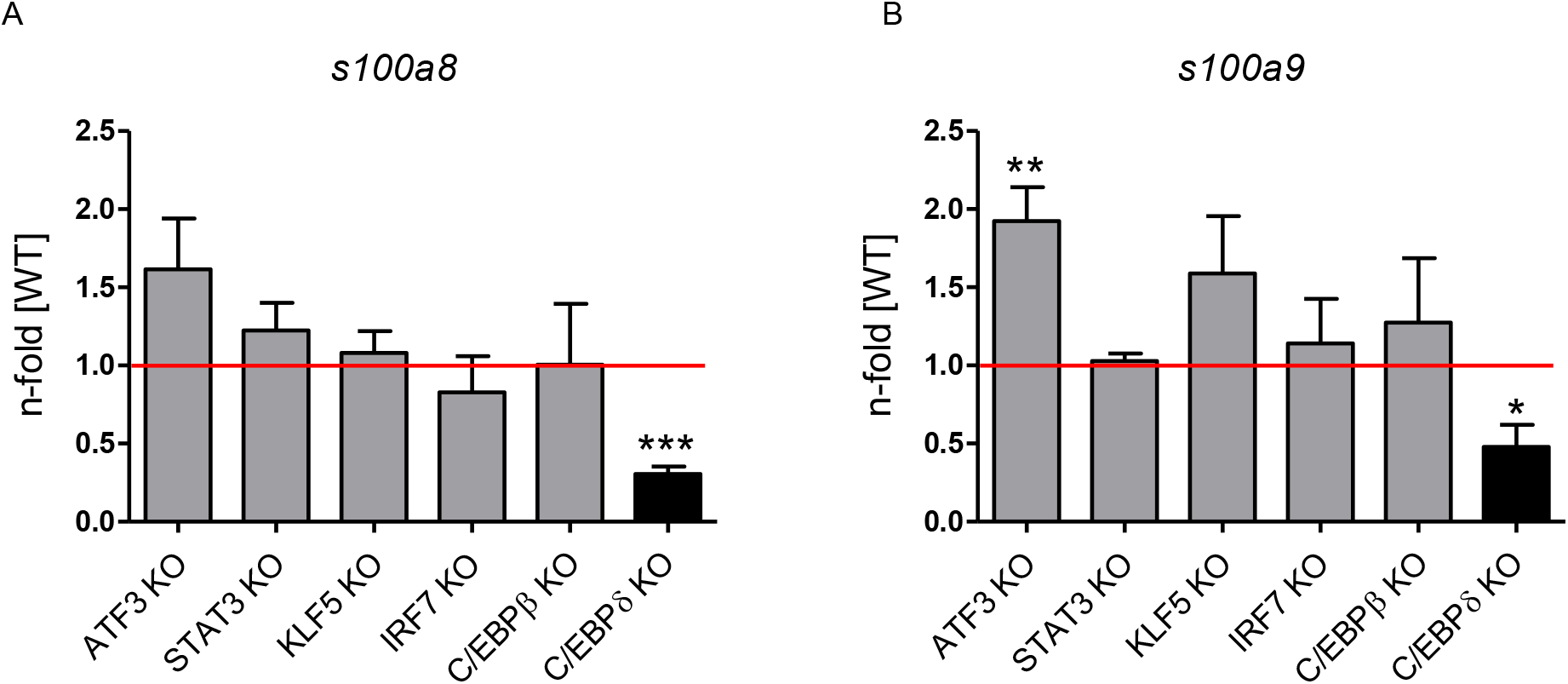
Relative S100A8 and S100A9 expression in differentiated single KO ERHoxb8 monocytes.

### TABLES

Table S1: Gene summary of MaGECK analysis (related to main Figure 1)

Table S2: List of guides (stated in 5’-3’ orientation) for cloning into lentiCRISPR v2, related to Methods

Table S3: List of primer (stated in 5’-3’ orientation) for amplifying GeCKO library and NGS, related to Methods

Table S4: List of oligonucleotides (stated in 5’-3’ orientation, fw: forward, rv: reverse) for cloning steps to construct TRE_3xFlag-C/EBPδ vector, related to Methods

Table S5: List of oligonucleotides (stated in 5’-3’ orientation, fw: forward, rv: reverse) for cloning steps to construct s100a8 and s100a9 reporter construct, related to Methods

Table S6: List of oligonucleotides (stated in 5’-3’ orientation) for mutagenesis to disrupt specific sites within s100a8 and s100a9 reporter vectors, related to Methods

Table S7: List of qRT-PCR primer (in 5’-3’ orientation) used for qRT-PCR, related to Methods

Table S8: List of ChIP-PCR primer (in 5’-3’ orientation) for s100a8 and s100a9 genomic locations, related to Methods

Table S9: Expression changes in the BioNRW monocytes dataset (RNA-seq, n=26) (related to main Figure 4)

